# A Super-Loop Extrusion Mechanism Shapes the 3D Mitotic Chromosome Folding

**DOI:** 10.1101/2025.05.05.652288

**Authors:** Davood Norouzi

## Abstract

Despite significant advances in understanding mitotic chromosome folding, existing loop-extrusion models – based on two nested tiers of Condensin I- and II-mediated loops – fail to fully reproduce the structural features of chromosomes observed in vivo. We propose a hierarchical three-layer model in which a third set of megabase-scale “super-loops,” extruded by a small number of high-processivity motors, compacts chromatin to its native dimensions. Polymer simulations of human chromosome 1 (250 Mb) show that 25-200 consecutive or interlaced super-extruders forming ∼10 Mb loops are sufficient to collapse a 25 μm bottle-brush structure into the empirically observed 5-8 × 1.5 μm rod. This model simultaneously reproduces the second diagonal in mitotic Hi-C maps, the double-helical scaffold seen in 3D-SIM microscopy, and the alternating-handedness, zero-mean helicity reported by high-resolution imaging. Interlaced super-loops enhance structural robustness against extruder loss and naturally account for the centromere as a region devoid of super-extruders. Although our simulations remain agnostic to molecular identity, we hypothesize that specialized forms of Condensin II, aided by auxiliary chromosome-associated proteins, act as super-extruders, while Topoisomerase II and other cross-linkers stabilize accumulated stress by converting transient DNA crossings into durable catenations. This three-layered mechanism resolves longstanding discrepancies in chromosome size, shape, contact-map profile, and scaffold ultrastructure within a unified physics-based framework. The next step is to identify or distinguish super-extruders experimentally and to design Hi-C, imaging, or mechanical perturbation assays to validate their existence and dynamics.

## Introduction

Our understanding of the structure and function of mitotic chromosomes has advanced significantly since these structures were first discovered over 140 years ago. However, many details remain unclear. During mitosis, chromatin undergoes extensive reorganization. It is well established that topoisomerase IIα, cohesin, chromokinesin KIF4A, and histone tail deacetylation are crucial for chromosome compaction, segregation, and maintenance [1-5]. However, the primary drivers of chromosome folding are the Condensin I and II structural maintenance of chromosomes (SMC) complexes [1-2]. These complexes have distinct yet complementary roles in organizing chromosomes into their characteristic rod-like shapes. Multiple studies have employed techniques such as Hi-C, live-cell imaging, acute depletion of SMC proteins with auxin degrons, and polymer modeling to investigate how these complexes restructure chromatin from its loose interphase organization into densely packed mitotic chromosomes [3-5].

Condensin II plays a key role early in mitosis, particularly during prophase, by initiating the formation of large chromatin loops through an ATP-driven DNA reeling mechanism within the nucleus. These loops, approximately 400-500 kb in size, are thought to be organized into a discontinuous helical scaffold, forming the primary structure of compacted chromatids [1-3]. The period length of this helical organization ranges from 8-15 Mb in mammals to 10-40 Mb in plants [4,6]. These periodic contacts are inferred from secondary diagonals seen in Hi-C interaction maps [4]. Hi-C (High-throughput Chromosome Conformation Capture with sequencing) is a genomic technique used to study the 3D structure of chromatin in the cell nucleus. It captures pairwise contact maps of DNA, revealing how DNA is spatially organized and how different regions of the genome interact [3].

Once the nuclear envelope breaks down in late prophase, Condensin I, which was previously confined to the cytoplasm, gains access to chromatin. It starts extruding smaller nested loops of around 80-100 kb within the larger formed loops. This hierarchical arrangement creates a dense “bottlebrush” structure. The combined action of Condensins I and II leads to chromosome compaction, producing the dense, rod-like morphology characteristic of metaphase chromosomes [1-2, 4]. Polymer modeling suggests that loop extrusion alone can compact DNA. Simulations indicate that the entropic forces arising from molecular volume exclusion between adjacent loops naturally produce rod-like chromosomes without requiring an external scaffold [4].

Despite the growing acceptance of the nested, two-layered loop-extrusion model, several questions remain. For instance, polymer models using consecutive Condensin I and II loops of ∼80 kb and ∼450 kb, respectively, result in overly long and thin chromosomes, which do not match actual mitotic structures. It has been proposed that Condensin II might adopt a spiral staircase arrangement, with loops extending in a helical pattern [4]. This model relies on fitting parameters such as the cylindrical structure’s radius and periodicity, with the latter derived from the second diagonal in Hi-C data. However, it remains unclear why Condensin II would naturally follow a helical arrangement. While some plant cells show clear helicity in chromosome structure [7], high-resolution 3D fluorescence imaging of mammalian cells reveals that mitotic chromosomes may not be helically coiled. Instead, they appear to have axes arranged as sequential half-helices of alternating handedness [8].

Advances in microscopy, such as serial block face scanning electron microscopy, allow whole-cell reconstruction of chromosomes at resolutions of 4 nm (cross-sectional) and 60 nm (axial), enabling precise measurements of chromosome length and volume. These studies indicate a linear relationship between chromosome volume/length and DNA length during metaphase and anaphase - relationships that current polymer models do not fully explain [5]. Additionally, recent findings using FIB/SEM electron microscopy and 3D-SIM reveal a double-helical scaffold within each chromatid. Note that what appears as a “double-helical scaffold” is an emergent artifact of the co-localization of non-histone proteins such as Condensin II and Topo IIα around the chromosome axis [9]. Other unresolved issues of the two-layered loop-extrusion model include the structural origin of the centromere, the role of Topo IIα, and the elastic properties of chromosomes that allow shape changes from prometaphase to anaphase.

### Proposed solution

Here, we propose a hierarchical model with three layers of loop extrusion instead of two. Layers one and two generate a cylindrical conformation that is too long and narrow, with Condensin I forming smaller loops nested within the larger Condensin II loops. Using polymer simulations, we show that the desired chromatin folding and shape can be achieved by introducing a third “super-extruder” that forms loops of size 3-15 Mb, depending on the cell type. These super loops can be either consecutive, interlaced or multi-laced, resulting in robust and stable chromosome structures. Consecutive super-extrusion can produce familiar shaped chromosomes, but these structures are easily disrupted. In contrast, interlaced super-extrusion is less sensitive to the exact placement and sizes of super loops and can tolerate instances where extruders fall off. Notably, very few super-extruders are needed. For instance, to compact human chromosome one, which is ∼250 Mb, only about 50 interlaced super-loops are required. The interlaced arrangement of super-extruders also appears to form two backbones, creating a double-helical scaffold. Additionally, this model of chromatin with super-loops predicts that centromeric regions simply lack super-extruders.

Our model reproduces the overall conformation of mitotic chromosomes but does not specify what super-extruders are, how they coordinate their orderly consecutive or interlaced arrangements, or how they determine loop size. Recent advances begin to fill these gaps: Hi-C and microscopy studies on single- and multiple-mutant degron cell lines now outline the rules that govern Condensin and cohesin engagement, traversal, stalling, and one-versus two-sided extrusion, clarifying how these activities shape sister-chromatid architecture [10]. A recent nanoscale DNA-tracing study shows that the rod shape of condensed chromosomes can arise without a contiguous protein scaffold; instead, Condensin traversals enable the formation of overlapping loops spanning several megabases, consistent with our super-loop extrusion hypothesis [11].

## Methods

We modeled each chromosome as a coarse-grained bead-spring polymer of 50000 beads, representing the length of human chromosome 1 with one bead corresponding to 5 kb of DNA. Mitotic Hi-C data and nucleosome density measurements support assigning a bead radius of r_0_ = 30 nm. Consecutive beads were linked by harmonic bonds and harmonic angles, while non-bonded interactions were handled with OpenMM’s [12] NonbondedForce using σ_vdW_ and ε_vdW_ values supplied in the input. To prevent unphysical self-energies during local bending and to maintain flexibility, we disabled non-bonded interactions for bead pairs separated by four or fewer monomers along the chain. Because the soft-sphere coarse-grained model has a maximum attainable density, we tested σ_vdW_ values, some smaller than the r_0_, ranging from 25 nm to 35 nm and observed only minor differences in chromatin compaction; σ_vdW_ = 28 nm was therefore adopted. The implementation allows each bead to carry its own parameter set, including electric charge, enabling monomer-specific variation. One such parameter is the designation of Condensin or super-extruder loop ends. These paired-bead interactions were applied with an OpenMM’s CustomBondForce defined by

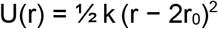

where k = 25 and 50 kJ mol^− 1^ nm^− 2^ for Condensin I and II loop ends respectively, and k = 80 kJ mol^− 1^ nm^− 2^ for super-extrusion looping. Simulations were performed on CUDA using a VariableLangevinIntegrator at 300 K with friction 0.003 ps^− 1^ and an integration error tolerance of 0.003. Following energy minimization (5000 steps, 1 kJ mol^− 1^ tolerance), dynamics were advanced in 5 × 10^5^-step blocks. The first 1 × 10^7^ ps ran without super-extruders, the next 4 × 10^7^ ps with super-extrusion active at ε_vdW_ = 0.1, and the final 5 × 10^7^ ps with ε_vdW_ = 0.3 to promote mild condensation. PDB and RST (restart) snapshots, together with per-force-group energies (bond, angle, torsion, non-bonded, custom), were recorded after every integration step block. Equilibration of the total energy is shown in Figure S1.

### Looping conformation initialization

Condensin II loops were generated first from a Gaussian distribution (mean = 480 kb, σ = 30 kb) with an average inter-loop gap of 50 kb and no overlaps. Condensin I loops were then randomly nested inside these larger loops, sampled from a Gaussian distribution (mean = 90 kb, σ = 10 kb) with a mean gap of 25 kb, also without overlap. Super-loops for human chromosome I were chosen from a normal distribution (mean = 9.7 Mb, σ = 0.4 Mb) to match the second-diagonal peak of Hi-C contact maps [4]. Three super-extrusion regimes were constructed: (**1**) consecutive, with non-overlapping loops separated by ∼60 kb gaps; (**2**) interlaced, where two loops overlap on average; and (**3**) multi-laced, allowing up to five overlapping loops at any position. We prepared 25 independent conformations for the consecutive regime, 25 for the interlaced regime, and 5 for the multi-laced regime, totaling 55 looping conformations for chromosome 1. All nested hierarchies of loops were resampled independently across these 55 conformations. Additional simulations were performed with super-loop sizes of 3 Mb, 5 Mb, 7 Mb, 12 Mb, and 15 Mb in both consecutive and interlaced configurations.

## Results

We first simulate two-layered nested loop formation, as shown in Figure 1a. The blue and yellow beads are the Condensin I and II loop ends pushed together. They produce a long, thin chromosome that is already cylindrical and rod-like without any scaffold or external constraints. But the structure is 25 μm long, which is not consistent with the expected size of less than 10 μm for human chromosome 1 [13].

**Figure 1:**
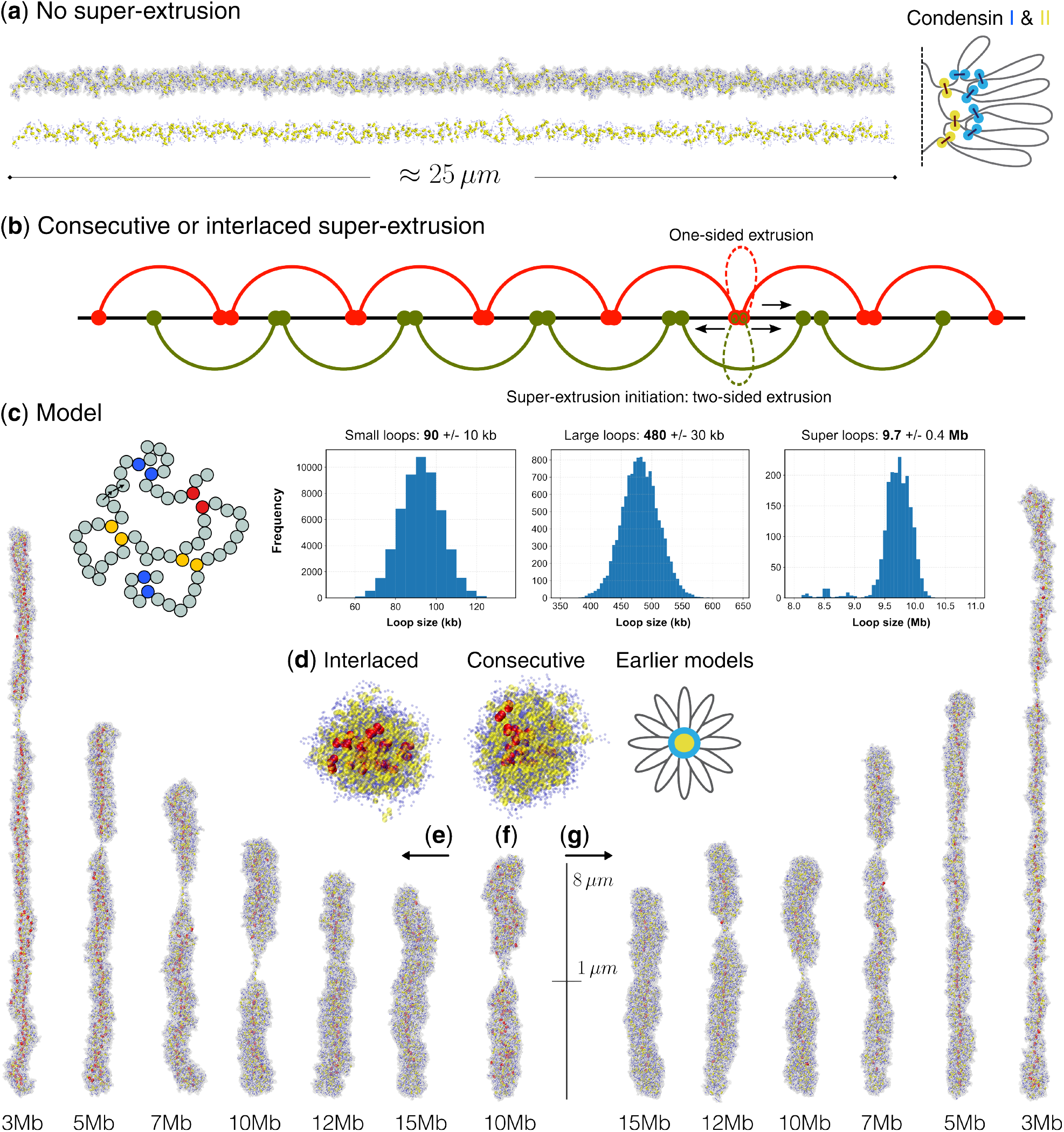
(**a**) Simulating 250 Mb of chromatin with Condensin I and II loop-generating SMC complexes results in a cylindrically stiff, rod-like conformation approximately 25 μm in length. (**b**) A set of super-loops is added to further condense the chromosome. These super-loops can be consecutive (non-overlapping) or interlaced. A combination of one-sided and/or two-sided loop extrusion can give rise to such looping regimes. (**c**) The coarse-grained DNA model represents chromatin as beads, with blue, yellow, and red beads indicating small-loop, large-loop, and super-loop ends, respectively. These ends are shown paired together. Histograms show the size distributions for small, large, and super-loops. (**d**) A top view of the simulated interlaced and consecutive conformations, contrasted with the earlier bottlebrush model of the chromosome. (**e**) Equilibrated interlaced structures with super-loop sizes ranging from 3-15 Mb, highlighting visible Condensins, super-extruders, and a clear centromeric region. (**f**) An equilibrated multilaced structure with super-loop size 10 Mb. (**g**) Equilibrated consecutive structures with super-loop sizes from 3-15 Mb, showing visible Condensins, super-extruders in red, and a defined centromeric region. The scale bar is 8 μm long and 1 μm wide.

This observation inspired us to consider a third layer of megabase-scale super-loops. We envisioned three types or regimes of super-extrusion, consecutive, interlaced and multi-laced (Figure 1b). The consecutive super-extrusion assumes super-loops are non-overlapping (red only loops in Figure 1b), while the other regimes assume dual or multi-laced overlapping super-loops (red+green loops in Figure 1b). The formation of consecutive super-extrusion can easily be managed by non-bypassing extruders, whereas interlaced super-extrusion could occur if two types of super-extruders initiate extrusion from the same locus - one performing one-sided extrusion and the other creating two-sided loops (Figure 1b). Multi-laced super-extruders are distributed randomly along the chromosome. While the details of super-extrusion initiation do not impact the results of the polymer simulations, it is important to hypothesize how these looping regimes arise.

The polymer-beads model of chromatin is depicted in Figure 1c, with blue, yellow, and red bead pairs pushed together to form the nested small, large, and super loops, respectively. Figure 1c also shows the loop-size distributions for the 55 simulations of human chromosome 1. Note that super-extruders are not permitted in the centromeric region (positions 122-125 Mb on human chromosome 1), which lies approximately halfway along the chromosome. In some simulations we place the centromere off-centre to model chromosomes with unequal arm lengths.

Snapshots of the chromosome are illustrated in Figure 1d, 1e, 1f, and 1g. Figure 1d shows the radial distribution of Condensin I, Condensin II, and super-extruders using a top view for consecutive and interlaced structures. While the super-extruders (red beads) are relatively centered, Condensin I and II are spread across the cross-section of the chromosome: Condensin I (blue beads) is distributed almost uniformly, and Condensin II (yellow beads) also covers most of the cross-section. This contrasts with earlier models of chromosome compaction, which assume Condensin II mostly accumulates at the centre with a halo of Condensin I around it [4].

Figure 1e, 1f, and 1g display sample snapshots of the chromosome under interlaced, multi-laced, and consecutive super-extrusion regimes. Sample structures simulated with super-extruders having average loop lengths of 3, 5, 7, 10, 12, and 15 Mb are shown. Larger super-loops compactify the chromosomes the most. A scale marker of length 8 μm and width 1 μm is provided as a ruler reference. Interlaced (1e), multi-laced (1f), and consecutive (1g) conformations with the same super-loop sizes are similar in overall chromosome length and width. DNA beads are rendered semi-transparent so that the positions of Condensins I, Condensins II, and super-extruders - in blue, yellow, and red, respectively - remain visible. The centromeric regions are visibly narrower than the rest of the chromosome. A chromosomal length of approximately 8 μm and width of 1.5 μm for conformations with super-loops of size 10-12 Mb appears compatible with human chromosome dimensions.

More detailed snapshots of simulated human chromosome 1, with an average super-loop length of 10 Mb, is provided in Figure 2. The interlaced super-loop architecture (Figure 2a) and the consecutive super-loop arrangement (Figure 2b) look identical on the surface, but the super-loops (red beads in the semi-transparent DNA model) reveal internal differences. The final equilibrated placement of super-extruder ends in space resembles an “emergent” backbone or scaffold, although distant super-loops are not in direct contact and are unaware of each other’s location. In consecutive, interlaced, and multi-laced structures, the emergent backbones resemble a relatively centered single spine, a dual helical ladder, and a more complex pattern, respectively (Figure 2c).

**Figure 2:**
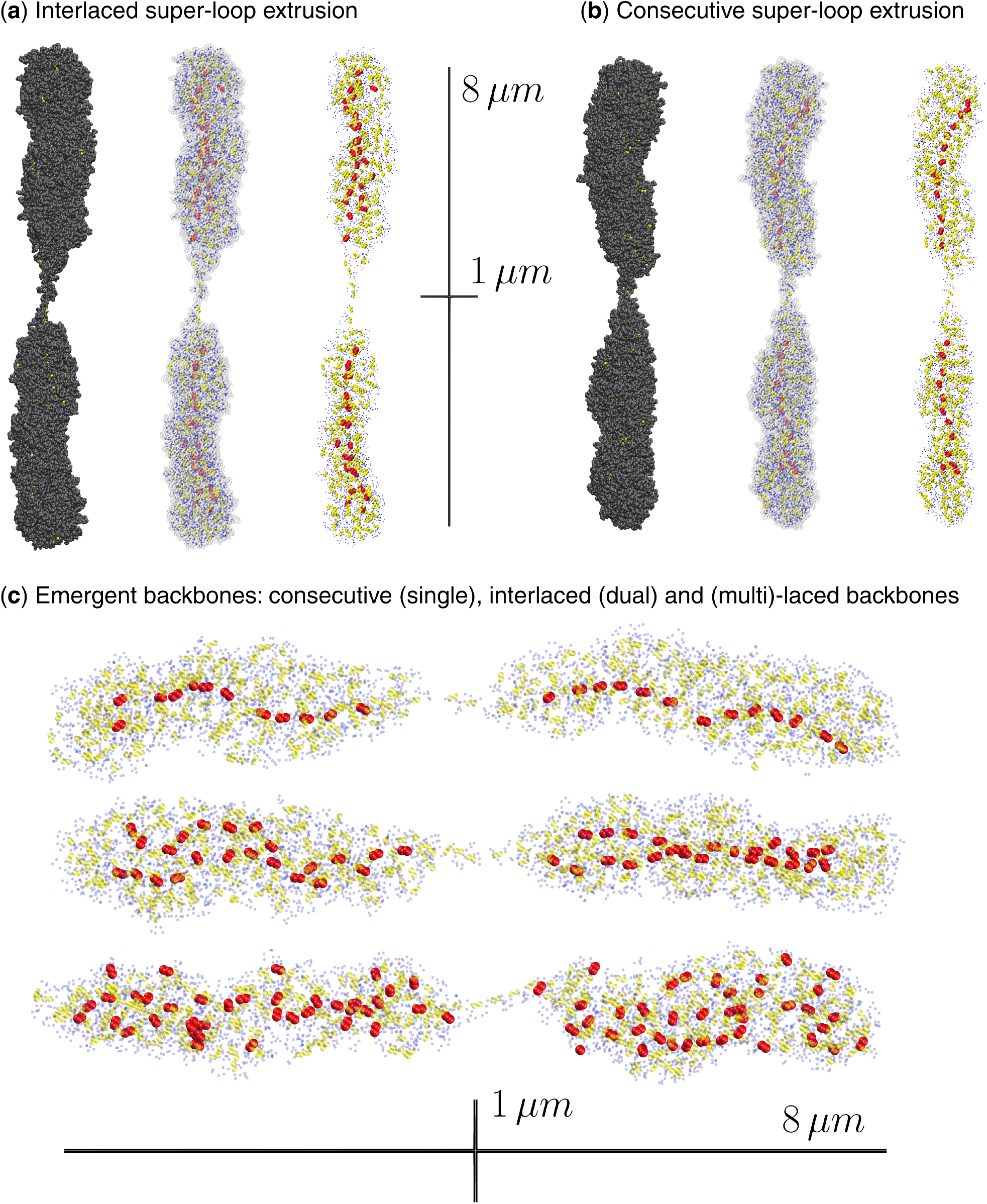
(**a**) A snapshot of a simulated human chromosome 1 with interlaced super-loops of size 10 Mb, shown in opaque, semi-transparent, and Condensins I (blue), II (yellow), and super-extruders (red) only views. The centromere region lacks super-extruders. The equilibrated structure is approximately 8 μm tall and 1.5 μm wide. (**b**) A similar snapshot is shown with three different views for the consecutive super-loop regime. (**c**) A closer look at the arrangement of super-extruder ends in the consecutive, interlaced, and multilaced regimes reveals emergent scaffolds resembling, respectively, a relatively centered single spine, a dual helical ladder, and a more complex pattern. The chromosome shape, size, and distribution of Condensin I and II appear unaffected by the choice (i.e., the number) of super-loops.

It is worth emphasizing that the idea of proposing and simulating interlaced super-loops originates from three-dimensional structured-illumination microscopy (3D-SIM) studies, which reported that the histone-depleted mitotic chromatin scaffold appears as a double-helical structure within each sister chromatid [9]. As previously mentioned, this arrangement also yields a more robust conformation, since the chromosome’s architecture remains intact even if some super-extruders fall off the chromosome.

Details of the simulation trajectories for interlaced and consecutive super-extrusion are provided as movies in the Supplementary Materials (Movie-S1 and Movie-S2) and are also available on YouTube (links below). The simulations begin with no super-extrusion, proceed with super-extruders activated, and then switch ε_vdW_ from 0.1 to 0.3 to simulate chromatin condensation induced by histone modifications or the effect of topoisomerase II or other cross-linkers. This change increases the packing density of the simulated chromosomes by approximately 30%.

Simulation trajectories after equilibration were used to compute statistical features of the folded chromosomes. In particular, we calculated contact frequency maps, which represent the probability that two monomers come into proximity as a function of their genomic separation. As expected, the likelihood of contact decreases with increasing genomic distance. Nonetheless, we observe prominent peaks corresponding to small, large, and super-loop sizes, consistent with the action of loop-extruding machinery bringing distant genomic regions into close contact. The resultant log-log contact frequency profiles are shown in Figure 3 and closely match experimental Hi-C data (Figure S2) [4], including the emergence and movement of the second diagonal peak towards ∼10 Mb as loop extrusion progresses. In experiments, Auxin-induced depletion of Condensin II causes the second diagonal in Hi-C maps to disappear [4], suggesting that super-extruders are likely Condensin II complexes equipped with specialized CAP (Chromosome-Associated Proteins) or other auxiliary subunits.

**Figure 3:**
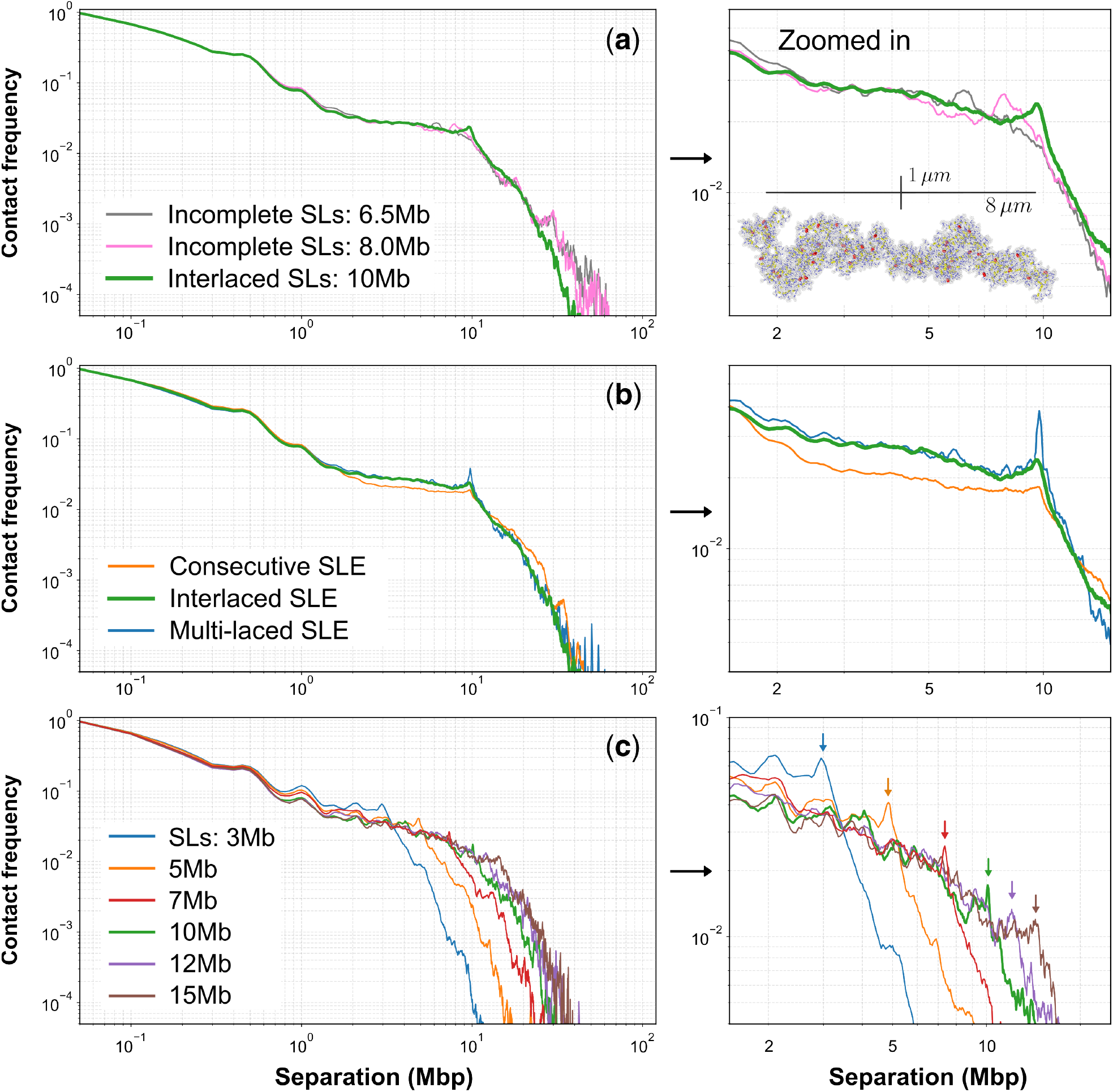
(**a**) The contact frequency profile - i.e., the probability that two monomers separated by a given genomic distance (in base pairs) are in close spatial contact - is shown. Monomers that are only a few beads apart along the DNA are more likely to contact than those near the chromosome ends. Due to forced looping mechanisms (e.g., small, large, and super-loop extrusion), peaks are expected at distances corresponding to loop sizes. Indeed, a prominent bump at ∼400–500 kb corresponds to Condensin II-mediated loops, and another peak at ∼10 Mb reflects the formation of super-loops. Contact frequency profiles for incomplete super-loop extrusion are also included. In the zoomed-in plot on the right, these peaks are more clearly resolved. A representative snapshot shows a chromosome in an intermediate stage of formation of super-loops. (**b**) Contact frequency profiles are shown for consecutive, interlaced, and multi-laced configurations. As the number of super-loops increases, the peak at ∼10 Mb becomes sharper. The zoomed-in plot on the right highlights these enhancements in detail. (**c**) Contact frequency profiles are plotted for configurations with super-loop sizes ranging from 3 to 15 Mb. Owing to the stiff, rod-like geometry of the chromosome, contact probabilities drop sharply beyond the super-loop size, since the polymer cannot bend sufficiently to bring distant monomers into contact. Clear peaks at 3, 5, 7, 10, 12, and 15 Mb are visible in the zoomed-in plot on the right.

Figure 3a shows a few noticeable bumps in the contact frequency profile, one at 400–500 kb distances and another at a separation distance of 10 Mb. Two additional contact frequency maps are also shown in Figure 3a for incomplete interlaced structures, where the super-loops are 6.5 and 8 Mb in size. Note that this differs from super-extrusion with similar loop sizes; i. e. in super-extrusion, the gap between super-loops is small, whereas in incomplete looping, the gap between super-loop ends spans several megabases. A zoomed-in plot of the contact frequency profile is provided in Figure 3a (right), which shows a gradual increase in the peak position as super-extrusion progresses. A sample snapshot of a chromosome during super-loop formation progression is shown within the plot.

Figure 3b and its zoomed-in version on the right compare super-loop extrusion (SLE) contact frequency profiles for consecutive, interlaced, and multi-laced regimes. As the number of super-loops increases through the addition of overlapping loops, more loci separated by ∼10 Mb are pulled toward each other via super-looping, causing the peak at this chromatin distance to become progressively sharper.

Figure 3c and its zoomed-in version on the right compare contact frequency profiles for simulations with super-loop (SL) sizes of 3, 5, 7, 10, 12, and 15 Mb. The sudden drop in proximity between distant monomers along the chromatin arises from the stiff, rod-like structure of chromosomes; i. e. beyond the super-loop distance (in base pairs), polymer beads rarely come into close spatial proximity in 3D space.

Interestingly, the super-extrusion model naturally produces a random helical structure in folded chromosome arms, with helicity emerging from buckling of the chromatin within each super-loop. Condensin I and II loops stiffen the chromatin (Figure 1a), and super-extruders applying further compaction to this stiffened structure can induce symmetry-breaking buckles; a well-known phenomenon in rod mechanics. When an elastic rod is compressed beyond a certain threshold, it bends in a random direction; for example, gently pressing the ends of a spaghetti strand causes it to buckle. A series of sterically hindered buckles formed by super-loops may result in a random helical conformation, with the overall helicity averaging close to zero in chromosomes with large super-loops.

To estimate the helicity, the ends of super-loops, used as reference points approximately equally spaced along the chromatin, were projected onto the chromosome’s cross-sectional (x–y) plane. The signed angle between adjacent reference points was calculated using the arccosine of the normalized dot product between neighboring endpoints. The cumulative sum of these angles provides a good estimate of how much the chromatin twists around the cylindrical (z-) axis.

Figures 4a and 4b illustrate this cumulative helical angle as a function of position along the chromosome (in degrees, with 360° corresponding to one full rotation) for interlaced and consecutive super-extrusion, respectively. The values shown are for single simulation snapshots and vary drastically between different equilibrated snapshots. The helicity changes sign frequently, and the total helicity at chromosome length 250 Mb generally remains close to zero in individual snapshots, and averages to zero across multiple, uncorrelated trajectories.

**Figure 4:**
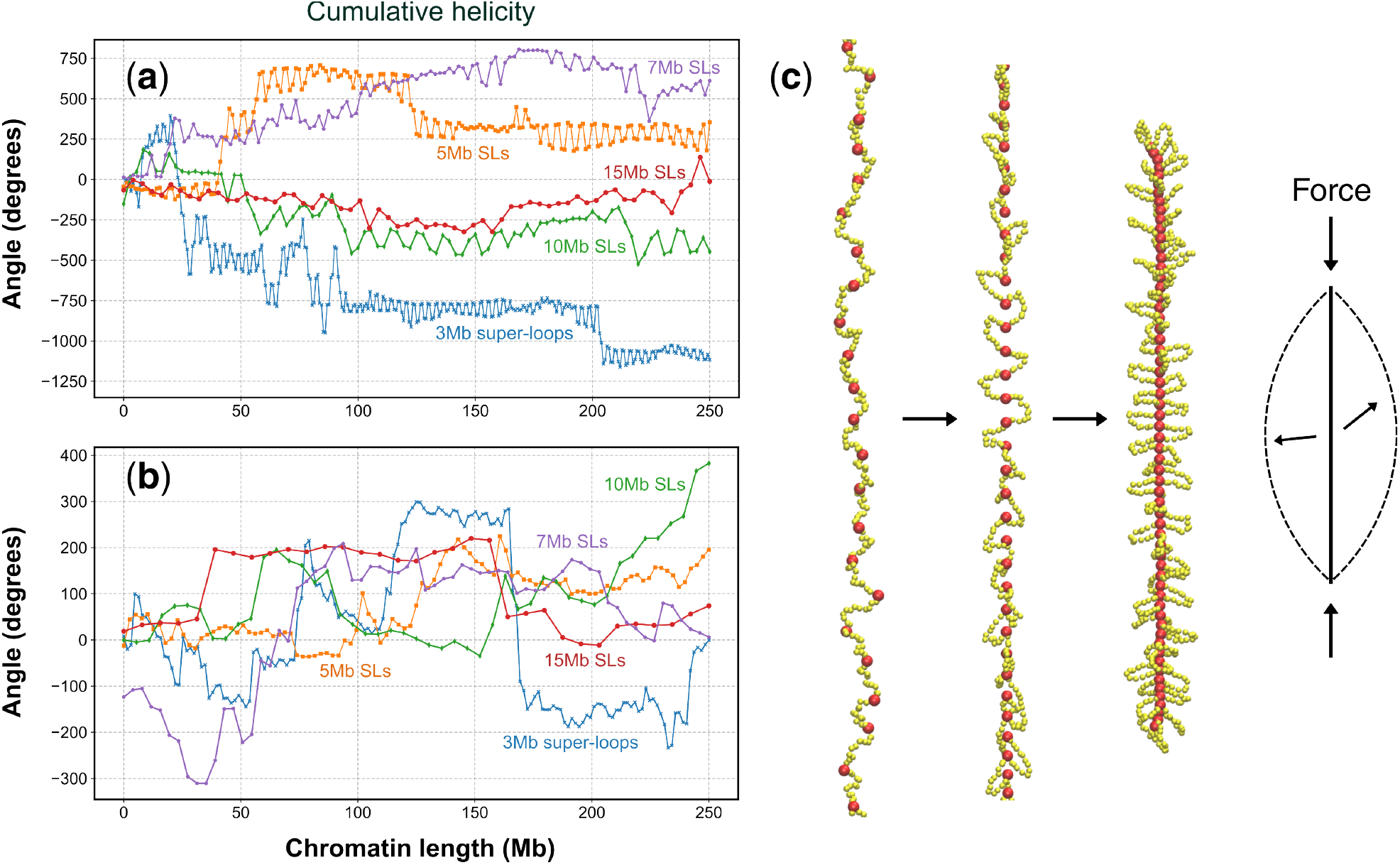
(**a**) Cumulative helicity is calculated by summing the signed angles between adjacent super-loops along the chromosome. This plot corresponds to interlaced configurations. The total winding number has reached -1100 degrees (about three full turns) in a 3 Mb super-loop conformation. However, taking an uncorrelated snapshot from the simulation can exhibit the opposite total helicity. Chromosomes with larger super-loops are less likely to exhibit significant overall twist. The cumulative winding displays local zigzag patterns, as adjacent interlaced loops alternate in helicity to avoid steric clashes. (**b**) Cumulative helicity for consecutive configurations is shown. These structures have more flexibility, and the overall twist tends to remain near 0 in all cases. (**c**) A combination of instantaneous symmetry-breaking buckling, driven by super-loop-extrusion pressure, and steric hindrance between buckled super-loops can result in locally helical regions with alternating handedness.

Figure 4c shows a trajectory of chromatin compaction, with super-loop ends in red and Condensin II endpoints in yellow. As super-extrusion progresses, internal Condensin II loops buckle away from super-loop ends coming into proximity. This buckling, combined with steric hindrance (not all DNA shown), creates local twists between adjacent super-loops. If these twists are in phase, the helicity accumulates to a net positive or negative twist; otherwise, the chromatin helicity appears as alternating half-helices with opposite handedness.

According to Figures 4a and 4b, the zigzag or alternating handedness is more apparent with interlaced super-extrusion. However, the structural constraints of the interlaced model allow helicity to grow larger, as it is less flexible in adjusting to local defects in helical twist. Nonetheless, even for interlaced chromatin, the total cumulative helicity remains close to zero and is greatest for smaller super-loops.

Super-loops of size 3-7 Mb impose more steric constraints on neighboring loops and can thus generate larger overall twists.

## Discussions

### Triple-layer loop extrusion resolves long-standing paradoxes in mitotic folding

Introducing a third, megabase-scale “super-extrusion” tier into the canonical Condensin I/II two-layer model converts an unrealistically long (25 µm) bottle-brush into a compact 8 × 1.5 µm rod that matches the observed dimensions of human chromosome 1. In this framework the centromere is simply a region devoid of super-loops. Depending on the looping regime, only ∼25, ∼50 or ∼200 super-extruders per chromatid - arranged consecutively, interlaced, or multilaced - are sufficient. Interlaced super-loops are especially robust. They tolerate lost extruders, recreate the double-helical scaffold seen by 3D-SIM and FIB/SEM, and reproduce the 10 Mb second diagonal in mitotic Hi-C. Our simulations also capture the experimentally observed stochastic helicity. The local buckling of the stiff Condensin I/II lattice under super-loop pressure produces alternating half-helices whose net twist averages to zero, consistent with high-resolution fluorescence reconstructions of mammalian chromosomes [8].

Contact frequency profiles derived from Hi-C data from prometaphase cells show that depletion of CAP-H, a key subunit of Condensin I, does not eliminate the second diagonal peak at 10 Mb. However, this peak disappears upon depletion of CAP-H2, which is specific to Condensin II. Since Condensin II forms large loops of approximately 400-500 kb, this suggests that it also plays a role in forming megabase-scale loops [4,11]. Whether additional auxiliary CAP subunits are required for super-loop formation remains to be experimentally verified.

### Topo II and auxiliary cross-linkers can lock in super-loop stress

Triple-layer extrusion concentrates mechanical load at super-loop bases. We propose that Topoisomerase IIα - assisted by linker proteins such as KIF4A, Ki-67, and histone H1 variants - locally relaxes twisted DNA while converting a subset of passage events into durable DNA catenations. These transient entanglements act as dynamic rivets, dissipating stress along the fiber and welding neighbouring extruder ends into the discontinuous scaffold visualised by EM staining. Although the average helicity over many snapshots is near zero, Topo II-dependent locking could freeze an individual chromosome in whatever twisted state happens to exist once super-loop extrusion ceases. This would make some chromosomes appear with substantial helicity in some species [7,16] especially those with smaller super-loop sizes ∼3-7Mb.

Auxin-induced depletion of Topo IIα has been shown to elongate chromosomes and reduce their width from approximately 1.4 µm to 0.6 µm -closely resembling our long, thin structures in the absence of super-loop extrusion (Figure 1a) [14]. This indirectly suggests that without Topo IIα-mediated entanglements, chromosome structure cannot be effectively stabilized.

### Why a 30 nm bead for 5 kb DNA is physically sound

Mitotic nucleosome density is estimated at 500-750 µMolar [5]. A sphere of radius 30 nm has a volume V≈1.1×10^−19^ liters. Packing ∼30 nucleosomes - the number in 5 kb of DNA - into that volume yields 30 / (N_A_ V) ≈ 450 µMolar, only modestly below the experimental range. For more compact chromosomes one could either reduce the bead radius or deepen the attractive well from the present ε_vdW_ = 0.3 kJ mol^− 1^ to 0.5 - 1 kJ mol^− 1^. Indeed, tests with r_0_ = 25 nm and ε_vdW_ = 0.75 shortened the simulated chromosome 1 to ∼5 µm tall. This degree of flexibility and compactness - i.e., a bead radius of 25 - 35 nm - is also consistent with flexibility and persistence length estimates from Monte Carlo simulations of nucleosomal arrays, which fit force spectroscopy measurements [15].

### Next steps

Future models should represent super-extruders explicitly as molecular motors that load, stall, and traverse DNA; incorporate centromeric stiffness gradients; and couple loop dynamics to histone-tail modifications to capture the length changes observed from prometaphase to anaphase. It is also worth exploring how the loop extrusion machinery precisely controls loop sizes. The relatively sharp second diagonal observed in most mitotic Hi-C maps suggests that loop sizes are tightly regulated. Notably, the 400-500 kb bump we observe in the contact frequency profiles - corresponding to Condensin II-mediated loops - is typically absent in experimental data [4]. This discrepancy may arise because, at such genomic scales, chromatin is enriched with cross-linking and other protein-mediated cis-interactions that obscure large-loop signals. Additionally, consecutive super-loops generate only a weak peak at the 10 Mb genomic separation, supporting the idea that super-loops in vivo are more likely to be interlaced or multi-laced.

Our computational modeling offers a promising solution to the long-standing puzzle of mitotic chromosome organization. Experimentally identifying the predicted ∼20-200 super-extruders per chromosome, however, remains a major challenge. If validated, this prediction would mark a significant breakthrough in using physics-based modeling to explain chromosome structure.

## Code Availability

https://github.com/norouzis/Super-loop-extrusion

## Acknowledgments

Grateful to all members of the Job Dekker lab for their support and feedback. We thank HHMI and UMass Chan Medical School. We also acknowledge the computational resources provided by the NIH, which enabled the simulations in this study.

## Supplementary Materials

Movie-S1: https://youtu.be/KXbXvvZTkDI Movie-S2: https://youtu.be/WdKiQTg3tKs

**Figure S1:**
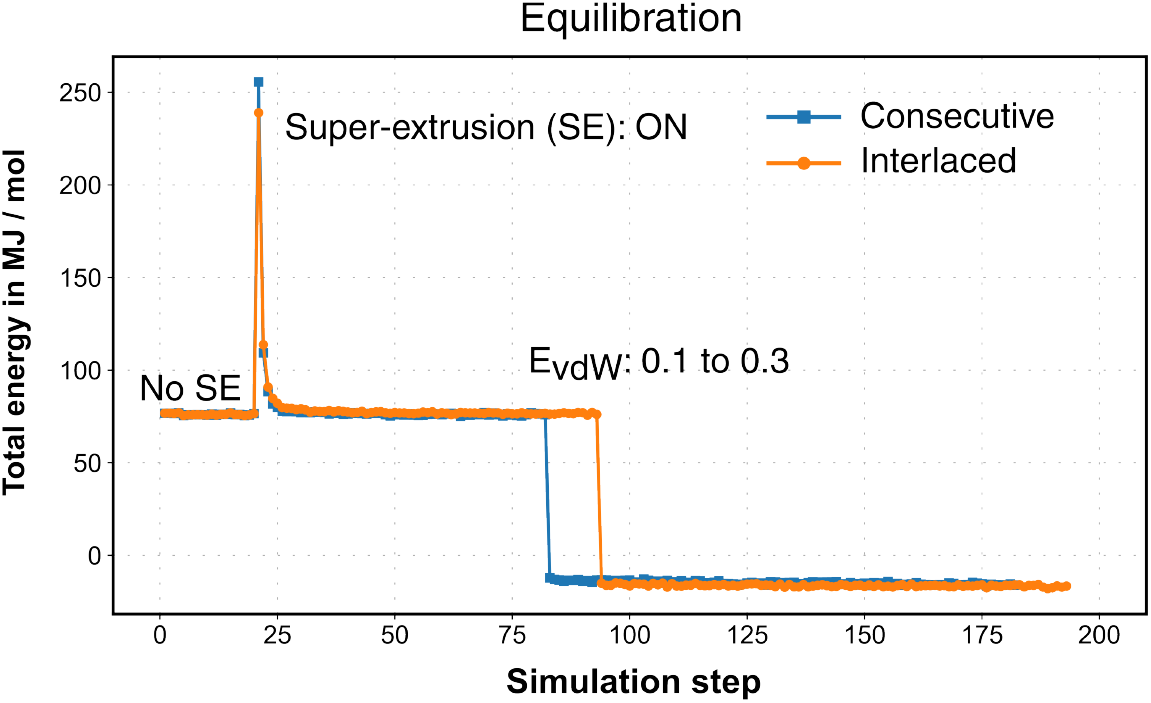
Equilibration was verified by monitoring the convergence of total energy. Representative simulation trajectories are selected. Initially, only Condensin I and II are active in the temporal region labeled “No SE.” Super-extrusion is then activated, causing the custom bond energy between distant super-loop ends to spike due to their initial separation. As compaction proceeds, the total energy decreases. To promote further condensation resembling phase separation, ε_vdW_ is increased from 0.1 to 0.3. Multiple conformations are sampled from the equilibrated structures for statistical analysis.

**Figure S2:**
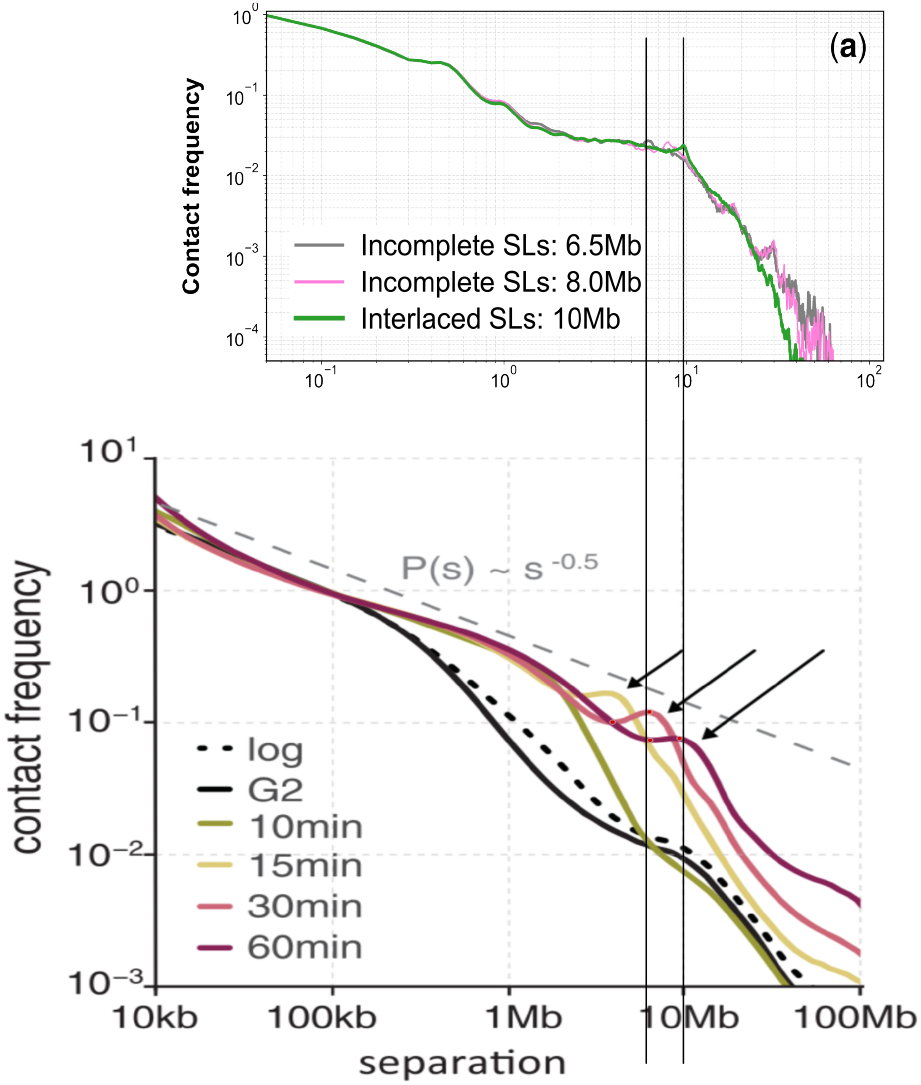
The contact frequency profiles of our simulated incomplete and completed interlaced super-extrusion structures match the peaks observed in Hi-C data [4]. A prominent peak around ∼10 Mb, along with the progressive emergence of the second diagonal in Hi-C maps, likely reflects the ongoing formation of super-loops.

